# NGAL drives cardiac dysfunction and fibrosis in rats with chronic kidney disease

**DOI:** 10.1101/2025.08.27.672761

**Authors:** M. Soulié, T. Sánchez-Bayuela, I. Lima-Posada, Y. Stephan, L. Nicol, Z. Lamiral, J. Lagrange, A. A. Voors, N. Lopez-Andres, N. Girerd, P. Mulder, F. Jaisser

## Abstract

**Background:** Patients with chronic kidney disease (CKD) are at high risk of cardiovascular (CV) complications. Neutrophil gelatinase-associated lipocalin (NGAL) is a well established marker of kidney injury, but recent evidence suggests that NGAL might also play an active in the progression of the cardiorenal syndrome.

**Methods:** CKD was induced in rats via 5/6 nephrectomy in wild-type (WT) and Ngal-knockout (KO) rats. Cardiorenal functions were assessed three months after subtotal nephrectomy or sham operation. Cardiac fibroblasts (CFs) were incubated with or without recombinant Ngal and galectin-3 (Gal-3).

**Results:** Cardiac function, including diastolic hemodynamics and perfusion, was less impaired in CKD Ngal KO than in CKD WT. Cardiac fibrosis was more severe in CKD WT than sham, but was blunted in CKD Ngal KO rats. Levels of Gal-3, collagen I, MCP-1 and IL-6 were elevated in cardiac fibroblasts incubated with recombinant Ngal. A similar pattern was observed in cells treated with recombinant Gal-3. Both Ngal and Gal-3 induced activation of the Tlr4-Myd88 pathway. Using Gal-3 or Tlr4 inhibitors, we showed that Gal-3 contributes to Ngal-induced cardiac fibrosis and inflammation by activating the Tlr4-Myd88 pathway. In patients with heart failure with preserved ejection fraction (HFpEF) (MEDIA-DHF and BIOSTAT-CHF cohorts), elevated levels of NGAL and Gal-3 were associated with pulmonary artery systolic pressure, a marker of advanced diastolic dysfunction and adverse clinical outcomes, particularly among patients with impaired renal function.

**Conclusion:** In non-diabetic CKD rats, Ngal was involved in the progression of diastolic dysfunction via a Gal-3/Tlr4-dependent pathway increasing inflammation and fibrosis.

## Introduction

Chronic kidney disease (CKD) is characterized by a progressive deterioration of renal structure and function. Cardiac and renal functions are closely intertwined, as impaired kidney function exacerbates heart failure and vice versa. CKD leads to volume overload and hypertension, increasing cardiac workload, promoting hypertrophy, and ultimately resulting in heart failure with preserved ejection fraction (HFpEF), a frequent complication in CKD patients^1^.

The renin-angiotensin-aldosterone system (RAAS) is activated in CKD, with angiotensin II and aldosterone (aldo) promoting cardiac inflammation and fibrosis^2–4^. High aldo levels are associated with a higher CV risk and higher risks of CKD progression and HFpEF development^5–7^ and higher risk of CKD progression^8^. Patients with high aldo levels also display greater deleterious LV remodeling^9^.

Treatment optimization is crucial to prevent CKD progression, reduce CV risks, and improve the quality of life of CKD patients. Targeting the aldo pathway, in addition to RAS blockade, has proved effective for reducing both CKD progression and CV risk in type 2 diabetic patients with renal failure^10^. Newer non-steroidal mineralocorticoid receptor (MR) antagonists, such as finerenone, prevent the progression of HFpEF in various patients, including those with impaired renal function^11,12^. However, the exact mechanisms of these beneficial effects in CKD remain unclear.

Moreover, the fear of hyperkalemia associated with aldo/MR blockade in CKD patients — although better controlled with non-steroidal MR antagonists such as finerenone — remains a barrier to the widespread use of MR antagonists. We previously identified neutrophil gelatinase-associated lipocalin (Ngal) as a downstream target of the aldo/MR pathway, demonstrating its essential role in the development of inflammation and fibrosis upon aldo/MR activation in the kidney and CV system^13,14^. These findings identify a potential novel therapeutic target within the MR pathway not associated with a risk of hyperkalemia. We investigated in the present study the potential of Ngal inhibition for preventing the development of CKD-associated HFpEF and the mechanisms involved.

## Materials and methods

The data generated and/or analyzed during this study are available from the corresponding author on reasonable request. A detailed methods section is provided in the supplemental material.

### Experimental design

Experiments were approved by the Darwin ethics committee of Sorbonne University (#22207-2019010411586046). They were conducted according to INSERM animal care guidelines in in accordance with DIRECTIVE 2010/63/EU of the European Parliament. Animals were housed in a facility with a controlled climate and a 12-h light/12-h dark cycle, with free access to water and food.

CKD was induced by 5/6 subtotal nephrectomy in 10-week-old Sprague Dawley rats (Charles River): two-thirds of the right kidney were removed in a first intervention, with the left kidney removed 10 days later. Ngal KO and WT littermates underwent the procedure simultaneously. In sham-operated rats, both kidneys were exposed, without resection. Ten days after the second intervention, plasma creatinine and urea determinations were performed to assess renal function. The generation of the Ngal KO mutant line is described in the supplemental methods, and a schematic diagram of the *in vivo* experimental design is provided in Fig. S1A.

### Study populations: MEDIA-DHF cohort, BIOSTAT-CHF cohort

The MEDIA-DHF cohort (Metabolic Road to Diastolic Heart Failure; NCT02446327) was a multinational, multicenter (10 centers), observational study that enrolled 626 patients. Eligible patients had (i) acute decompensated HF, (ii) a recent (within the last 60 days) hospital discharge for acute HF, or (iii) chronic stable HF in an ambulatory setting. The primary endpoint was the composite of cardiovascular death (CVD) or first HF hospitalization (HFH) within a 12-month period of follow-up.^15,16^ The BIOSTAT-CHF study (A systems BIOlogy Study to TAilored Treatment in Chronic Heart Failure; EudraCT 2010-020808-29) was a prospective, multicenter study conducted across 11 European countries, enrolling adult patients with new-onset or worsening HF, defined on the basis of symptoms and either a left ventricular ejection fraction (LVEF) ≤40% or high levels of natriuretic peptides (BNP >400 pg/mL or NT-proBNP >2000 pg/mL). The primary outcome was the composite of all-cause mortality and rehospitalization for HF within a two-year period of follow-up. Each component was also analyzed separately as a secondary outcome. For the analysis presented here, we considered only patients with HFpEF^17^. Both studies adhered to the principles of the Declaration of Helsinki and were approved by the competent ethics committees. All participants gave written informed consent.

### Statistical analysis

The results are presented as the mean ± SEM. Differences in the means between groups were assessed by two-way repeated-measures ANOVA (protocol 2) comparing four groups, provided that the data were normally distributed for each group (Shapiro-Wilk normality test, in GraphPad Prism 8.0.1). Analyses were performed with GraphPad Prism 8.0.1 and differences were considered significant if *p* <0.05.

For clinical studies, statistical analyses were performed with SAS version 9.4. We considered *p*-values <0.05 in two-tailed tests to indicate statistical significance for main effects, whereas *p*<0.10 were considered significant for interaction terms. Results are presented as beta coefficients for linear models and hazard ratios for survival analyses, each with 95% confidence intervals (95% CI). In the MEDIA-DHF cohort, linear regression models were used to evaluate the association between pulmonary arterial systolic pressure (PASP) and biomarker levels (NGAL or galectin-3, BIOTECHNE, Minneapolis, USA), while accounting for renal function status (eGFR <60 vs. ≥60 mL/min/1.73 m²) and potential interactions between biomarkers and renal function. Analyses were stratified by renal function and performed with three levels of adjustment (univariable adjustment and adjustment for age and sex) and fully adjusted models including adjustment for age, sex, body mass index, diabetes, and hypertension. The assumptions underlying linear regression were systematically verified. Cox proportional hazards models were used in both MEDIA-DHF and BIOSTAT-CHF to investigate the associations between biomarker levels (NGAL and GAL-3) and clinical outcomes, namely cardiovascular death or CV hospitalization in the MEDIA cohort, and all-cause death and HF hospitalization in the BIOSTAT study. Interactions were assessed in models incorporating the biomarker concerned, renal function category, and their multiplicative interaction. The proportional hazards assumption was tested with covariates varying over time. For each outcome, we report crude, age- and sex-adjusted, and fully adjusted hazard ratios.

## Results

### CKD induces CV alterations after one month in both WT and KO Ngal rats

We evaluated the renal and CV consequences of CKD induction by 5/6 nephrectomy. Renal dysfunction was similar in CKD Ngal KO and CKD WT rats, with high plasma urea and creatinine levels, albuminuria or low GFR. Renal fibrosis was also similar in the two strains upon CKD induction. (Fig. S2A-G).

Cardiac Ngal levels were higher in CKD WT rats than in the sham-operated WT group, whereas Ngal was undetectable in the sham-operated and CKD Ngal KO groups, as expected, indicating efficient *lcn2* gene inactivation (Fig. S1B). Cardiac echocardiography parameters in the CKD WT and KO groups were similar to those in the corresponding sham-operated groups, with no significant difference between in CKD WT and CKD Ngal KO rats (Table S1). Fractional shortening was lower in CKD Ngal rats than in Ngal Sham-operated rats but not in CKD WT rats relative to sham-operated WT rats (Table S1). The cardiac output of CKD rats was similar to that in the corresponding sham-operated rats. (Table S1). LV and body weights were similar in all groups (Table S1).

Systolic blood pressure (SBP) and diastolic blood pressure (DBP) were not affected in the sham-operated or CKD rats of either the WT or Ngal KO groups (Table S1). Neither LV end systolic pressure nor LVESPVR was affected by CKD in WT or Ngal KO rats relative to the corresponding sham-operated animals (Table S1). LVEDP was higher in WT and Ngal KO rats than in the corresponding sham-operated animals and did not differ significantly between CKD Ngal KO and CKD WT rats (Fig. 1A). LVEDPVR, an index of diastolic dysfunction, was higher in CKD WT rats than in sham-operated controls. This increase was blunted in CKD Ngal KO rats relative to CKD WT rats, revealing a protective effect of the absence of Ngal expression on diastolic function (Fig. 1B).

**Figure 1.**
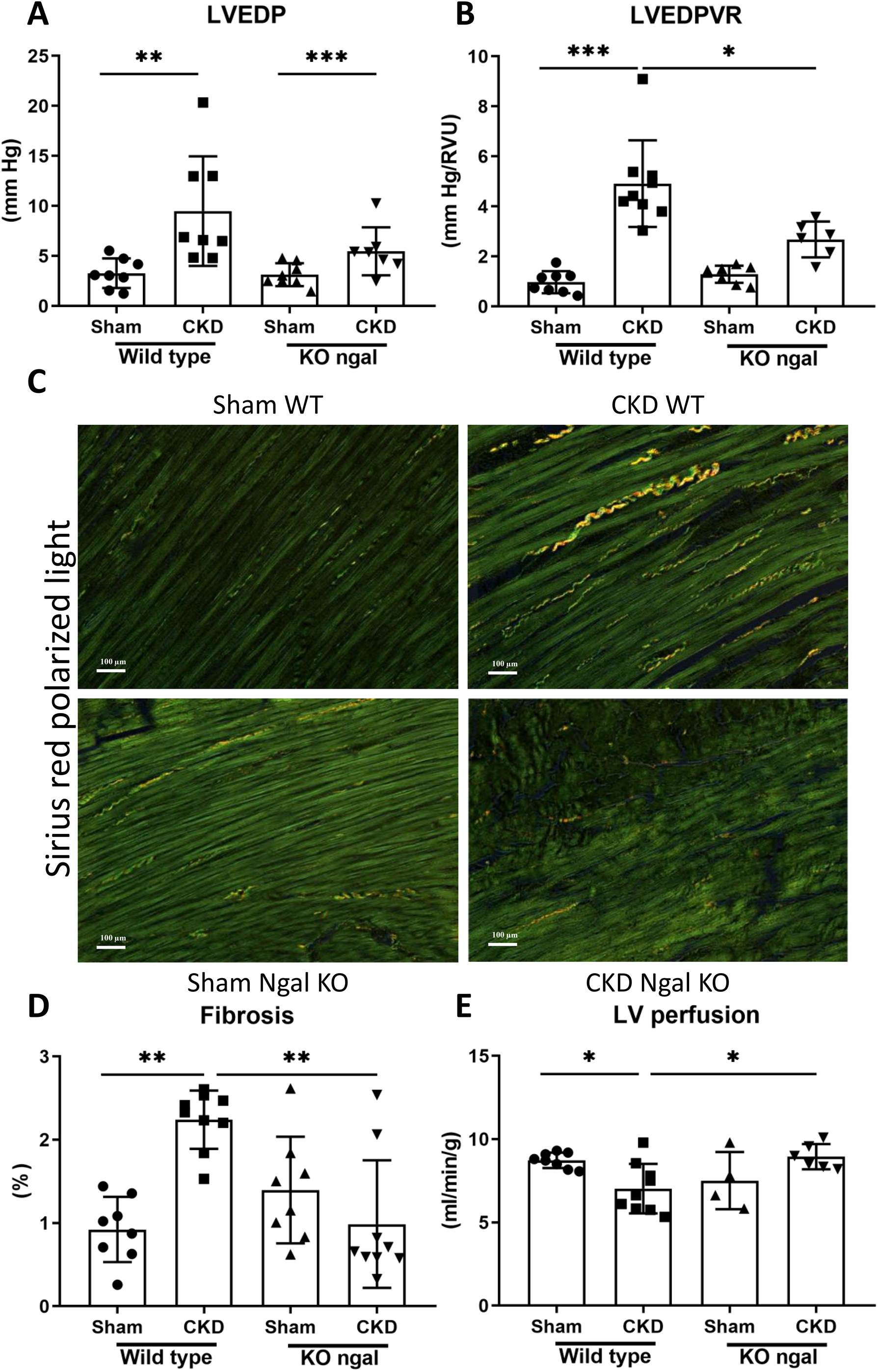
Ngal knockout improves cardiac function and fibrosis in rats with chronic kidney disease. CKD: chronic kidney disease. **A** Left ventricle end diastolic pressure (LVEDP, mmHg), **B** left ventricle end diastolic pressure volume relationship (LVEDPVR, mm Hg/RVU), **C** cardiac fibrosis (illustration) **D** fibrosis (%). **E** Myocardial perfusion (mL/min/g). The data are presented as the mean ± SEM, *n* = 8–10. Statistical analysis was performed by one-way ANOVA followed by Tukey’s post hoc test. \**p* < 0.05, \*\**p* < 0.01, \*\*\**p* < 0.001.

Interstitial cardiac fibrosis was increased in CKD WT rats compared to sham-operated WT rats (Fig. 1C-D). However, the increase in cardiac fibrosis was less marked in CKD Ngal KO rats than in CKD WT rats. (Fig. 1C-D). LV cardiac perfusion was impaired in CKD WT rats relative to sham-operated WT rats (Fig. 1E). LV cardiac perfusion in CKD Ngal KO rats was greater than that in CKD WT rats (Fig. 1E).

### Aldosterone induces the expression of profibrotic and inflammatory genes, including *Ngal* and *Gal3*, in rat CFs, an effect abolished by cotreatment with finerenone

We previously reported that aldo increases the levels of profibrotic and proinflammatory markers in human CFs, an effect blunted by the non-steroidal MR antagonist finerenone^13^. We obtained similar results in rat cardiac fibroblasts, making it possible to perform further mechanistic studies in rat CFs. Relative levels of fibrosis and inflammation marker gene expression (*Col1A1*, *Col3A1*, *Ctgf*, *Ccl2*, *CT 1*, *IL-6)* were increased by aldo in rat CFs, an effect abolished by finerenone (Fig. S3A-D). Expression of the *Ngal* and *Gal-3* genes was also increased by aldo treatment in rat CFs, an effect blunted by finerenone (Fig. S3E-F).

### Recombinant Ngal and Gal-3 induce fibrotic and inflammatory markers in CFs

Recombinant Ngal (rNgal) increased the secretion of Ccl2 and IL-6 into the supernatant of CFs after 24 h of exposure and the secretion of Col1 and Gal-3 after 48 hours of exposure (Fig. 2A). Similar effects were observed with recombinant Gal-3 (rGal-3), which induced increases in the levels of Ccl2, IL-6 and Col1 secretion after 24 h or 48 h (Fig. 2B). The expression of genes involved in fibrosis and inflammation, such as *Col1A1*, *Col3A1*, *Ccl2* and *IL-6,* was increased by both rNgal (Fig. 2C) and rGal-3 (Fig. 2D). As expected, co-incubation with the Gal-3 inhibitor MCP blunted the increase in gene expression observed after co-incubation with rGal-3 (Fig 2D). Importantly co-incubation with MCP, a Gal-3 inhibitor, also blunted the effects of rNgal, highlighting the crucial role of Gal-3 in the profibrotic and proinflammatory effects of Ngal (Fig. 2C).

**Figure 2.**
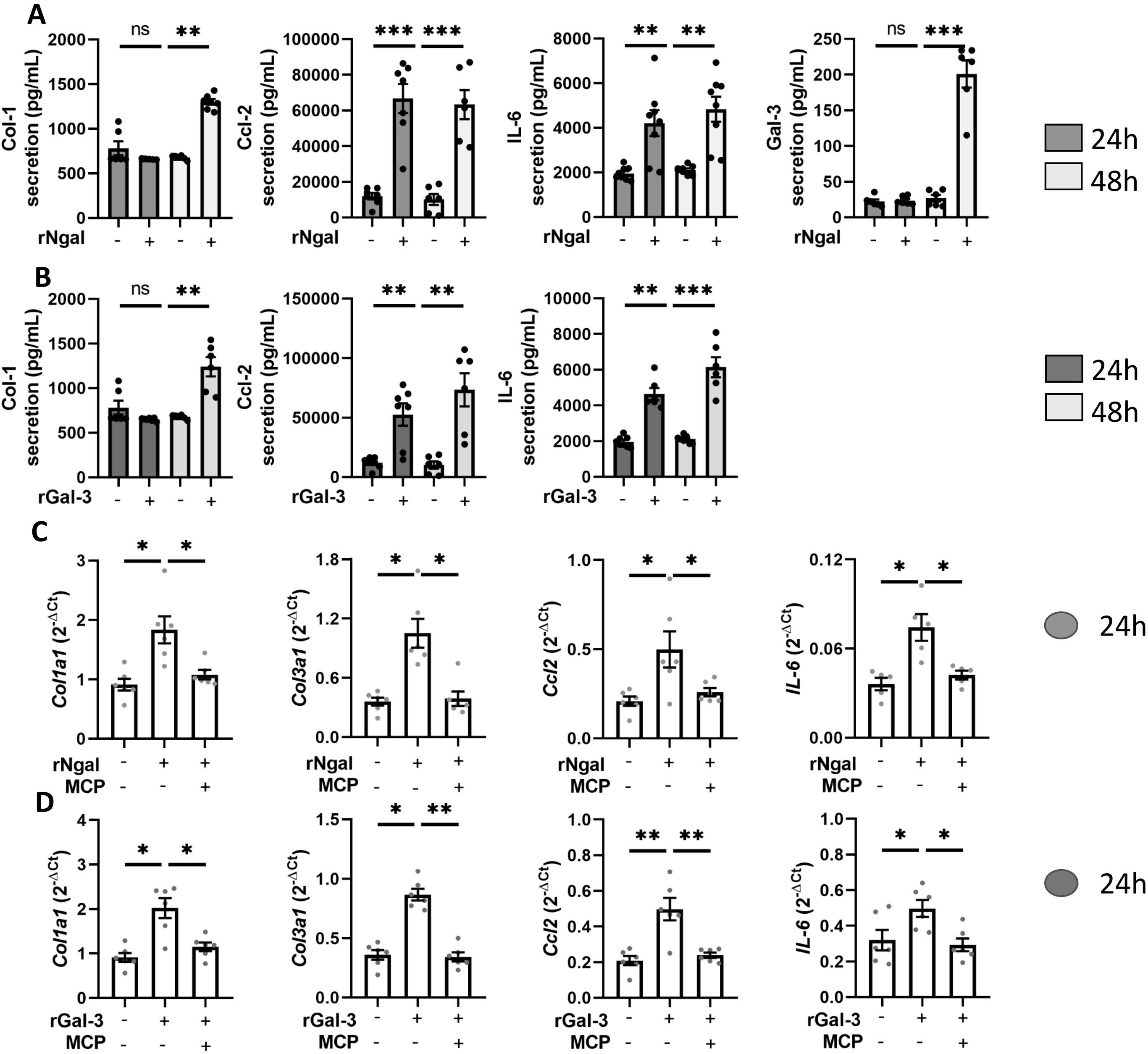
Recombinant Ngal and Galectin-3 induce fibrotic and inflammatory proteins in a Gal-3 pathway-dependent manner in cardiac fibroblasts. **A** Col1 (pg/mL), Ccl2 (pg/mL), IL-6 (pg/mL) and galectin-3 (Gal-3) secretion by cardiac fibroblasts (CFs) incubated with rNga;, **B** Col1 (pg/mL), Ccl 2 (pg/mL) and IL-6 (pg/mL) secretion by CFs incubated with rGal-3; **C** Expression (ΔCt values) of *Col1a1*, *Col3a1*, *Ccl 2* and *IL-6* genes in CFs incubated with rNgal with or without modified citrus pectin (MCP), a Gal-3 inhibitor. **D**. Expression (ΔCt values) of *Col1a1*, *Col3a1*, *Ccl 2* and *IL-6* genes in CFs incubated with rGal-3 with or without modified citrus pectin (MCP), a Gal-3 inhibitor. Data are presented as the mean ± SEM, *n* = 5–6 biological replicates. Statistical analysis was performed by one-way ANOVA followed by Tukey’s post hoc test. \**p* < 0.05, \*\**p* < 0.01, \*\*\**p* < 0.001.

### The profibrotic and proinflammatory effects of Ngal and Gal-3 in CFs are mediated by the Tlr4-Myd88 innate immunity pathway

Both rNgal and rGal-3 increased the expression, at both mRNA and protein levels, of the innate immune response genes *Tlr4* and *Myd88* (Fig. 3A-B) and the expression of genes involved in downstream Tlr4 signaling, such as *Irak1* and *Irak4* (Fig. 3B-C). The effects of rNgal and rGal-3 on Tlr4 pathway activation were inhibited by the Gal-3 inhibitor MCP (Fig. 3B-C). As expected, induction of the *Tlr4, Myd88, Irak1* and *Irak4* genes was blunted by the cotreatment with TAK-242 (a Tlr4 inhibitor) of CFs treated with rNgal or rGal-3 (Fig. S4A). Inhibition of the Tlr4 pathway with TAK-242 also blunted the increases in Col1, Ccl2 and IL-6 secretion (Fig. 4A-B) and the increases in *Col1A1*, *Col3A3*, *Ccl 2* and *IL-6* gene expression (Fig. S4B-C) induced by both rNgal and rGal-3, demonstrating the crucial role of the Tlr4-Myd88 pathway in Ngal-Gal-3 signal transduction resulting in profibrotic and proinflammatory effects.

**Figure 3.**
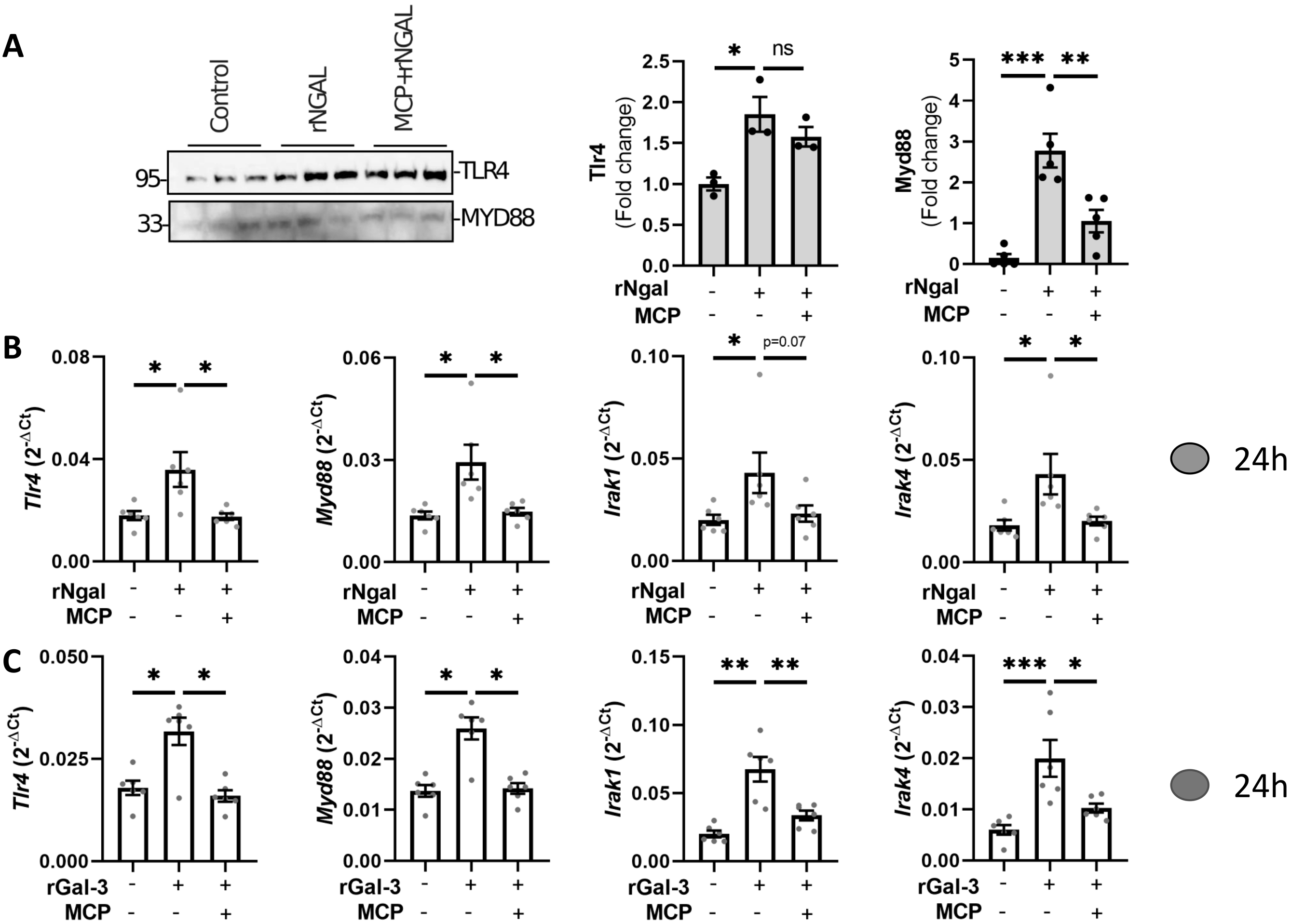
Galectin-3-dependent activation of Tlr4/Myd88 signaling by recombinant Ngal and Galectin-3. **A** Tlr4 and Myd88 protein levels in CFs incubated with rNgal with or without modified citrus pectin (MCP), a galectin-3 (Gal-3) inhibitor (left panel, representative image of the western blot; right panel, relative expression levels), **B** Expression (ΔCt values) of *Tlr4*, *Myd88*, *Irak1* and *Irak4* in CFs incubated with rNgal with or without modified citrus pectin (MCP), a galectin-3 (Gal-3) inhibitor, in cardiac fibroblast (CFs). **C** Expression (ΔCt values) of *Tlr4*, *Myd88*, *Irak1* and *Irak4* in CFs incubated with rGal-3 with or without modified citrus pectin (MCP), a galectin-3 (Gal-3) inhibitor, in cardiac fibroblast (CFs). Data are presented as the mean ± SEM, *n* = 5–6 biological replicates. Statistical analysis was performed by one-way ANOVA followed by Tukey’s post hoc test. \**p* < 0.05, \*\**p* < 0.01, \*\*\**p* < 0.001.

**Figure 4.**
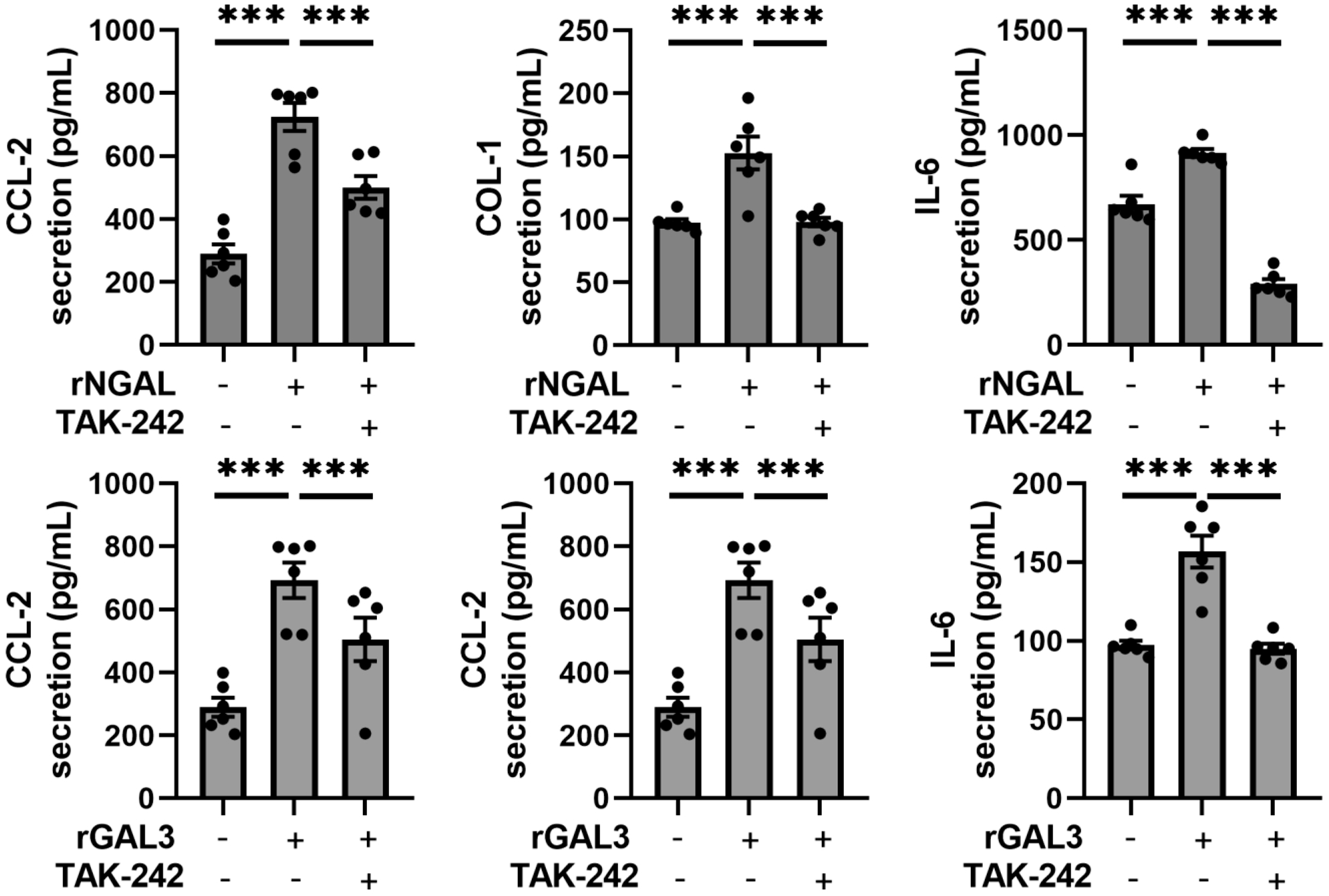
TLR4 signaling is required for rNgal- and rGal-3-driven inflammatory and fibrotic responses. **A** Col1 (pg/mL), Ccl2 (pg/mL) and IL-6 (pg/mL) secretion into the supernatant of cardiac fibroblasts (CFs) incubated for 24 h with rNgal with or without the Tlr4 inhibitor TAK-242; **B** Col1 (pg/mL), Ccl 2 (pg/mL) and IL-6 (pg/mL) secretion into the supernatant of CFs incubated for 24 h with rGal-3 with or without TAK-242. Data are presented as the mean ± SEM, *n* = 5–6 biological replicates. Statistical analysis was performed by one-way ANOVA followed by Tukey’s post hoc test. \**p* < 0.05, \*\**p* < 0.01, \*\*\**p* < 0.001.

### Correlation between NGAL, GAL-3 and CV outcomes in the MEDIA-DHF and BIOSTAT-HF cohorts

In patients with HFpEF from the MEDIA-DHF and BIOSTAT-HF cohorts, NGAL and GAL-3 levels were moderately correlated. In MEDIA-DHF, the Spearman correlation coefficient for the relationship between NGAL and GAL-3 (Olink) was 0.27 (0.19–0.36, *p*<0.0001). In BIOSTAT-HF, NGAL was more strongly correlated with GAL-3 determinations by mass spectrometry (r=0.53) than with those obtained with Olink (r=0.16) (both *p*<0.0001), suggesting platform-dependent variation.

On stratification for renal function (eGFR <60 vs. ≥60 mL/min/1.73 m²), both biomarkers were found to display differential association with PASP, a marker of advanced diastolic dysfunction (Table 1). In MEDIA-DHF, NGAL was significantly associated with PASP only in patients with low eGFR values (β=3.08 mmHg, 0.93–5.24, *p*=0.005), with no association found in patients with preserved renal function (β=–0.16 mmHg, *p*=0.95). Similarly, high GAL-3 levels were associated with a higher PASP only in patients with an eGFR <60 (β=6.76 mmHg, 1.95–11.57, *p*=0.006), not in those with an eGFR ≥60 (β=3.28 mmHg, *p*=0.15).

**Table 1.** Association of NGAL/GAL-3 levels with PASP in patients with HFpEF from the MEDIA-DHF cohort according to renal function (<60 or ≥60 mL/min/1.73 m^2^) In the Metabolic Road to Diastolic Heart Failure (MEDIA-DHF) cohort, the associations between pulmonary artery systolic pressure (PASP) and the biomarkers NGAL and galectin-3 (GAL-3) are presented as beta coefficients with their corresponding 95% confidence intervals (CI). Analyses were stratified by renal function (eGFR: estimated glomerular filtration rate) and performed with three levels of adjustment: univariable, age- and sex-adjusted, and fully adjusted models including adjustment for age, sex, body mass index, diabetes, and hypertension. Cox proportional hazards models were used to investigate associations with the levels of the NGAL and GAL-3 biomarkers. For main effects, statistical significance was defined as *p*-values <0.05 in two-tailed tests (*), whereas *p* < 0.10 was considered to indicate significance in assessments of interaction terms.

| MEDIA-DHF cohort |  | eGFR <60<br>N=146 |  | eGFR ≥60<br>N=179 |  |  |
| --- | --- | --- | --- | --- | --- | --- |
| Variable | Model | Beta (95 % CI) | P-value | Beta (95 % CI) | P-value | P-interaction |
| NGAL (ng/mL) | Univariable | 2.72 (0.54 ; 4.89) | 0.01 | 0.29 (-4.48 ; 5.06) | 0.91 | 0.36 |
|  | Sex age adjusted | 2.84 (0.70 ; 4.97) | 0.009 | -0.22 (-4.90 ; 4.45) | 0.93 | 0.24 |
|  | Full adjustment | 3.08 (0.93 ; 5.24) | 0.005 | -0.16 (-4.85 ; 4.53) | 0.95 | 0.22 |
| GAL-3 (per log-2 unit) | Univariable | 5.93 (1.06 ; 10.81) | 0.02 | 3.25 (-1.31 ; 7.81) | 0.16 | 0.43 |
|  | Sex age adjusted | 6.81 (2.00 ; 11.62) | 0.006 | 3.59 (-0.90 ; 8.09) | 0.12 | 0.33 |
|  | Full adjustment | 6.76 (1.95 ; 11.57) | 0.006 | 3.28 (-1.24 ; 7.80) | 0.15 | 0.29 |

A similar pattern was observed for clinical outcomes (Table 1). In MEDIA-DHF, high GAL-3 levels were associated with a higher risk of cardiovascular hospitalization or death in the eGFR <60 group (HR=2.18, 0.95–5.00, *p*=0.07 age- and sex-adjusted; HR=2.21, 0.95–5.15, *p*=0.07 fully adjusted), but not in the eGFR ≥60 group (HR=0.72 and 0.73, both *p*>0.5), with an interaction of borderline significance (*p*=0.09). NGAL was not significantly associated with outcomes regardless of renal function.

In BIOSTAT-HF, NGAL was associated with all-cause death (ACD) in patients with an eGFR <60 (HR=1.60, 1.16–2.19, *p*=0.004), but not in those with an eGFR ≥60 (HR=0.63, *p*=0.37), with an interaction of borderline significance (*p*=0.09 and 0.08 in age- and sex-adjusted and fully adjusted models, respectively). GAL-3 levels determined by mass spectrometry followed a similar pattern: significant association with ACD in the eGFR <60 group (HR=1.27 per 10 ng/mL, *p*=0.004) with no significant association in the group with preserved eGFR (*p*=0.72), with an interaction p=0.04 (age- and sex-adjusted) and 0.08 (fully adjusted). Similar results were obtained for GAL-3 levels measured by proximity extension assay (HR=1.61, *p*=0.03 for eGFR <60; HR=0.90, *p*=0.72 for eGFR ≥60), with an interaction, *p*=0.04 and 0.11, respectively. No differential pattern was observed when HFH was considered (all *p* values for interaction >0.20; table 2).

**Table 2:** Association of NGAL and GAL-3 levels with clinical outcome according to renal function in patients with HFpEF from the MEDIA-DHF and BIOSTAT-HF cohorts. Analyses were conducted within the Metabolic Road to Diastolic Heart Failure (MEDIA-DHF) and the BIOlogy Study to TAilored Treatment in Chronic Heart Failure (BIOSTAT-HF) cohorts. The associations of NGAL and galectin-3 (GAL-3) levels with cardiovascular hospitalization (CVH), cardiovascular death (CVD), all-cause death (ACD), and hospitalization for heart failure (HFH) were evaluated. Results are presented as beta coefficients for linear models and hazard ratios (HR) for survival analyses, together with 95% confidence intervals (CI). Analyses were stratified by renal function (eGFR: estimated glomerular filtration rate) and performed with three levels of adjustment: univariable, age- and sex-adjusted, and fully adjusted models including adjustment for age, sex, body mass index, diabetes, and hypertension. Cox proportional hazards models were used to investigate associations with levels of the biomarkers NGAL and GAL-3. For main effects, statistical significance was defined as *p*-values <0.05 in two-tailed tests (*), whereas *p* < 0.10 was considered to indicate significance in assessments of interaction terms.

| MEDIA-DHF cohort |  |  | eGFR <60<br>N=190 |  | eGFR >=60<br>N=236 |  |  |
| --- | --- | --- | --- | --- | --- | --- | --- |
| Variable | Ouctome | Model | HR (95%CI) | P-value | HR (95%CI) | P-value | P interaction |
| NGAL (per<br>ng/mL) | CVH/CVD | Univariable | 1.02 (0.73 ; 1.43) | 0.90 | 0.62 (0.22 ; 1.72) | 0.36 | 0.36 |
|  |  | Sex age<br>adjusted | 1.03 (0.73 ; 1.44) | 0.86 | 0.65 (0.24 ; 1.78) | 0.40 | 0.39 |
|  |  | Full<br>adjustment | 1.02 (0.72 ; 1.43) | 0.92 | 0.66 (0.24 ; 1.78) | 0.41 | 0.42 |
| GAL-3 (per<br>log-2 unit) | CVH/CVD | Univariable | **2.23 (0.97 ; 5.13)\*** | **0.06** | 0.74 (0.28 ; 1.99) | 0.55 | **0.09** |
|  |  | Sex age<br>adjusted | **2.18 (0.95 ; 4.96)\*** | **0.07** | 0.72 (0.27 ; 1.95) | 0.52 | **0.09** |
|  |  | Full<br>adjustment | **2.21 (0.95 ; 5.15)\*** | **0.07** | 0.73 (0.27 ; 1.98) | 0.53 | **0.09** |
| BIOSTAT-HF cohort |  |  | eGFR <60<br>N=1002 |  | eGFR >=60<br>N=1215 |  |  |
| Variable | Outcome | Model | HR (95%CI) | P-value | HR (95%CI) | P-value | P interaction |
| NGAL (per<br>ng/mL) | heart failure<br>hospitalization | Univariable | 1.15 (0.80 ; 1.64) | 0.45 | 1.28 (0.51 ; 3.19) | 0.60 | 0.83 |
|  |  | Sex age<br>adjusted | 1.14 (0.79 ; 1.65) | 0.48 | 1.09 (0.43 ; 2.75) | 0.86 | 0.92 |
|  |  | Full<br>adjustment | 1.17 (0.81 ; 1.67) | 0.41 | 1.0 (0.40 ; 2.51) | 1.00 | 0.77 |
|  | all-cause<br>death | Univariable | **1.48 (1.10 ; 2.00)\*** | **0.01** | 0.76 (0.29 ; 1.98) | 0.57 | 0.19 |
|  |  | Sex age<br>adjusted | **1.52 (1.13 ; 2.06)\*** | **0.007** | 0.61 (0.22 ; 1.68) | 0.34 | **0.09** |
|  |  | Full<br>adjustment | **1.60 (1.16 ; 2.19)\*** | **0.004** | 0.63 (0.23 ; 1.71) | 0.37 | **0.08** |
| GAL-3 (per<br>log-2 unit) | heart failure<br>hospitalization | Univariable | 0.98 (0.62 ; 1.53) | 0.91 | 1.41 (0.75 ; 2.62) | 0.28 | 0.35 |
|  |  | Sex age<br>adjusted | 0.96 (0.60 ; 1.53) | 0.86 | 1.45 (0.77 ; 2.75) | 0.25 | 0.31 |
|  |  | Full<br>adjustment | 0.94 (0.59 ; 1.51) | 0.80 | 1.47 (0.76 ; 2.83) | 0.25 | 0.29 |
|  | all-cause<br>death | Univariable | **1.60 (1.09 ; 2.33)\*** | **0.02** | 0.93 (0.55 ; 1.58) | 0.80 | 0.10 |
|  |  | Sex age<br>adjusted | **1.76 (1.17 ; 2.63)\*** | **0.006** | 0.87 (0.51 ; 1.49) | 0.62 | **0.04** |
|  |  | Full<br>adjustment | **1.61 (1.06 ; 2.45)\*** | **0.03** | 0.90 (0.52 ; 1.58) | 0.72 | 0.11 |
| GAL-3 (per<br>10 ng/mL) | heart failure<br>hospitalization | Univariable | 1.14 (0.94 ; 1.38) | 0.17 | **1.22 (1.02 ; 1.46)\*** | **0.03** | 0.62 |
|  |  | Sex age<br>adjusted | 1.16 (0.95 ; 1.42) | 0.15 | **1.21 (1.01 ; 1.46)\*** | **0.04** | 0.75 |
|  |  | Full<br>adjustment | 1.20 (0.97 ; 1.47) | 0.087 | **1.22 (1.01 ; 1.48)\*** | **0.047** | 0.89 |
|  | all-cause<br>death | Univariable | **1.21 (1.04 ; 1.41)\*** | **0.01** | 0.93 (0.73 ; 1.19) | 0.56 | **0.07** |
|  |  | Sex age<br>adjusted | **1.25 (1.06 ; 1.47)\*** | **0.007** | 0.91 (0.70 ; 1.17) | 0.46 | **0.04** |
|  |  | Full<br>adjustment | **1.27 (1.08 ; 1.49)\*** | **0.004** | 0.95 (0.74 ; 1.24) | 0.72 | **0.08** |

These findings, consistent across cohorts and assay platforms, suggest that NGAL is correlated with GAL-3 and that the prognostic value of NGAL and GAL-3 for HFpEF is modified by renal function.

## Discussion

Our findings indicate that Ngal inactivation has a cardioprotective effect in 5/6 nephrectomy rats. Genetic knockout of *lcn2* and an absence of Ngal are associated with milder diastolic dysfunction and fibrosis, and enhanced cardiac perfusion relative to WT littermates with CKD. This protective effect may be partially attributed to a lower level of CF activation. We observed an overexpression of profibrotic and inflammatory genes in CFs following incubation with aldo, Ngal, or Gal-3. This effect was attenuated by the MR antagonist finerenone and the inhibitors of Gal-3 and Tlr4. These results suggest a pathway in which MR overactivation leads to increases in Ngal production and Gal-3 levels, with downstream activation of the innate immune response through the Tlr4-Myd88 pathway

### Role of Ngal in the diastolic dysfunction and cardiac inflammation associated with CKD

Ngal, also known as Lcn2, is a 25 kDa protein from the lipocalin family involved in the innate immune response^18^. Ngal has emerged as a biomarker of renal injury and CV stress. It is considered an early biomarker of renal injury as its levels increase markedly within two to four hours of acute renal injury^18^. Plasma Ngal levels are also a good indicator of CKD development in patients^19^. Ngal levels have been associated with an increase in CV risk^20,21^. In the STANISLAS cohort, serum Ngal levels were positively correlated with systolic pressure and negatively correlated with sodium excretion levels^22^. High levels of Ngal in HF patients indicate underlying renal dysfunction and inflammation, which are closely linked to the progression of heart failure^23^.

The mechanisms underlying the deleterious effects of Ngal in the renal and CV systems remain unclear. We previously showed that Ngal expression is controlled directly by aldo via the MR and mediates MR-induced inflammation and fibrosis in various organs (kidney, heart, vessels) and cell types (renal distal tubule cells, immune cells, including macrophages in particular, and cardiac fibroblasts)^24^. In the immune system, Ngal favors the M1 polarization of macrophages by upregulating the response to IFNγ and LPS and suppressing IL-10 signaling^25^. Ngal deficiency has been shown to decrease the recruitment of neutrophils and macrophages in the heart after ischemia/reperfusion^25^. We have shown that Ngal is essential for cardiac remodeling and dysfunction after myocardial infarction, via the NF-κB pathway^13^. We have also shown that Ngal plays a crucial role in aldo-induced renal fibrosis and inflammation, by modulating IL-4 signaling in renal macrophages^26^. Together, these results highlight the potentially detrimental role of Ngal in the cardiac remodeling associated with CKD, as observed here in CKD Ngal KO rats.

Excess collagen deposition by activated CFs plays a crucial role in increasing cardiac stiffness, disrupting normal heart compliance through the secretion of collagen (types I, III, and VI) and fibronectin, the main components of the extracellular matrix (ECM)^27^. This process results in myocardial stiffening and impaired diastolic function, especially in patients with HFpEF, in whom cardiac stiffness is directly dependent on collagen^28^. In chronic remodeling scenarios, such as hypertension or post-infarction, sustained fibroblast activation is marked by persistent ECM overproduction^29^. We therefore focused our mechanistic studies on CFs to understand the pathways linking Ngal and cardiac remodeling.

### Gal-3, a target of Ngal, is essential for Ngal signaling in rat CFs

Ngal modulated the expression of multiple profibrotic and proinflammatory markers in CFs. We focused on Gal-3, a β-galactoside-binding lectin that we had previously identified as a key mediator of mineralocorticoid-induced cardiac remodeling^30^. Gal-3 is secreted by activated macrophages and plays a central role in myocardial fibrotic and inflammatory remodeling by promoting myofibroblast proliferation and collagen deposition^31^. Gal-3 has also emerged as a clinically relevant biomarker in CKD, in which high plasma Gal-3 levels are associated with disease progression^32^. Studies of patients with HFpEF have reported a correlation between Gal-3 levels and specific diastolic function parameters, such as E/e′ ratio, LV mass, and functional performance metrics, including peak VO₂ and 6-minute walk distance^33^. Clinical studies have also reinforced links between plasma Gal-3 concentration and new-onset HFpEF, poor prognosis, and the severity of cardiac structural and functional alterations in HFpEF populations^34,35^. These findings suggest that Gal-3 promotes inflammation and myocardial fibrosis, thereby contributing to diastolic dysfunction and adverse cardiac remodeling^35^. Importantly, the pharmacological inhibition of Gal-3 with MCP attenuated the profibrotic and proinflammatory responses induced by recombinant Ngal in CFs, suggesting that Gal-3 acts downstream from Ngal to exert its deleterious fibrotic and inflammatory effects.

### Ngal induces the Tlr4-Myd88 pathway via Gal-3 in CFs

The Tlr4-MyD88 signaling pathway plays a key role in linking innate immune activation to cardiac dysfunction. Upon recognition of pathogen-associated or damage-associated molecular patterns, Tlr4 engages the adaptor protein MyD88, initiating downstream signaling cascades that result in the activation of nuclear factor-kappa B (NF-κB) and the production of proinflammatory cytokines^36^. This inflammatory environment promotes CF activation and the deposition of extracellular matrix components, which in turn impair myocardial compliance^37^. In mice, deletion of the *myd88* gene or the pharmacological inhibition of Myd88 results in lower levels of fibrosis in the lung, kidney or liver. Preclinical studies have shown that deletion of the *Tlr4* or *Myd88* gene in mice attenuates fibrosis and improves diastolic function in angiotensin II-induced hypertension^38^. In the context of CKD, sustained activation of the Tlr4-Myd88 axis in cardiac tissue may therefore contribute to adverse cardiac remodeling, including hypertrophy and fibrosis. We show here that Ngal induces Gal-3, thereby increasing Tlr4-Myd88 signaling in CFs. The blockade of Tlr4-Myd88 signaling with the Tlr4 antagonist TAK-242 showed that this pathway was essential for the profibrotic and proinflammatory effects of Ngal and its downstream target, Gal-3, in CFs.

### Clinical relevance

The clinical findings reported here are consistent with the NGAL/GAL-3 mechanisms demonstrated in our experimental models. In both settings, high NGAL and GAL-3 levels were clearly associated with high pulmonary artery systolic pressure, a hallmark of advanced diastolic dysfunction, and with adverse outcomes, particularly in patients with impaired renal function. These human data provide a translational validation of our preclinical findings, highlighting the critical role of renal function in modulating the relationship between the NGAL/GAL-3 axis and the severity and prognosis of HFpEF. Supporting evidence from other clinical studies further reinforces this link. In patients with diabetes mellitus, plasma NGAL levels are independently associated with LV hypertrophy, even after adjusting for age, sex, BMI, eGFR, and SBP^39^. A recent meta-analysis also found that plasma GAL-3 serves as a predictive biomarker for new-onset HFpEF, adverse prognosis in established HFpEF, and the severity of LV diastolic dysfunction. Notably, in patients with HF with reduced ejection fraction, NGAL levels are independently correlated with GAL-3 levels, even after adjusting for potential confounders such as age, ejection fraction, and eGFR^34^.

This underscores the therapeutic potential of targeting this pathway, especially in patients with concomitant HFpEF and renal dysfunction, a subgroup at markedly high risk for poor clinical outcomes.

## Conclusion

Our results show that inactivation of the *ngal* gene in rats is cardioprotective in the context of CKD. We found that Ngal activates fibrotic and inflammatory pathways via Gal-3 and the Tlr4-Myd88 axis in CFs. Targeting Ngal is therefore a potential novel therapeutic approach for limiting HFpEF development in CKD.

## Data Availability Statement

The data generated and/or analyzed during the current study are available from the corresponding author upon reasonable request.

## Acknowledgments

We thank the Institut Clinique de la Souris – PHENOMIN-ICS for expert assistance in establishing the lcn2/ Ngal mutant rat line. We are also grateful for the excellent technical assistance and support for animal care provided by the *Centre d’Exploration Fonctionelles*.

## Author Contributions

F.J. and N.A.L. provided the concept and design of research M.S., T.S.B., I.L.P., L.N., Y.S., and P.M. performed experiments and prepared figures, M.S., T.S.B., P.M., N. L-A., N.G. and F.J. analyzed data and interpreted the results of experiments. F.J., N.G., A.A.V., J., L. and M.S., drafted the manuscript. M.S., T.S.B., I.L.P., Y.S., L.N., P.M., N.A.L. N.G. and F.J. approved the final version of the manuscript.

## Perspectives

NGAL has emerged as a biomarker of renal injury. Our data suggest that targeting Ngal may be of interest for limiting HFpEF development in CKD. NGAL may therefore represent a novel therapeutic target in this and potentially other contexts.

## Novelty and significance

### What is new?

- Rat Ngal gene inactivation is cardioprotective in the context of CKD
- We describe a novel Ngal signaling pathway via gal-3-Tlr4 that could be targeted in CKD-associated CV dysfunction.
- High NGAL/GAL-3 levels in patients are predictive of a poorer cardiac outcome, particularly when associated with eGFR<60 mL/min/1.73m 2

### Significance

The development of novel therapeutic approaches to target organ damage — including extracellular matrix remodeling and inflammation, which are common underlying mechanisms of cardiac dysfunction associated with renal disease — will be a major goal over the next decade. Neutrophil gelatinase-associated lipocalin modulates the inflammation, fibrosis and, ultimately, cardiac damage associated with CKD.

## List of nonstandard abbreviations and acronyms

ACD: All-cause death
Aldo: Aldosterone
Ccl2/Mcp1: C-C motif chemokine ligand 2 or monocyte chemoattractant protein 1
CFs: Cardiac fibroblast
CI: Confidence interval
Col1: Collagen 1
*Col1a1*: Collagen type I alpha 1 chain
*Col3a1*: Collagen type III alpha 1 chain
CVH: Cardiovascular hospitalization
Gal-3: Galectin-3
LVEDPVR: Left ventricle end diastolic pressure volume relationship
MCP: Modified citrus pectin
MR: Mineralocorticoid receptor
Myd88: Myeloid differentiation primary response 88
Ngal: Neutrophil gelatinase-associated lipocalin (Lcn2)
PASP: pulmonary arterial systolic pressure
rGAL-3: Recombinant galectin 3
rNgal: Recombinant neutrophil gelatinase-associated lipocalin
TAK-242: Toll-like receptor 4 inhibitor
Tlr4: Toll-like receptor 4

